# PDCD1 expression increases at elevated temperatures

**DOI:** 10.1101/2025.05.22.652424

**Authors:** Agnieszka Toma-Jonik, Patryk Janus, Katarzyna Mrowiec, Natalia Vydra, Kinga Sarkowicz, Monika Bar, Justyna Mirek, Magdalena Olbryt, Wojciech Fidyk, Wiesława Widłak

**Affiliations:** Center for Translational Research and Molecular Biology of Cancer, Maria Skłodowska-Curie National Research Institute of Oncology, Gliwice Branch, 44-102 Gliwice, Poland; Department of Bone Marrow Transplantation and Oncohematology, Maria Skłodowska-Curie National Research Institute of Oncology, Gliwice Branch, 44-102 Gliwice, Poland

**Author notes:** **Corresponding authors:** Agnieszka Toma-Jonik, Wiesława Widłak. These authors contributed equally to this work (Agnieszka Toma-Jonik, Patryk Janus).

**Keywords:** fever, heat shock (HS), HSF1, hyperthermia, immune checkpoints (ICP)

## Abstract

PDCD1 (Programmed cell death protein 1) is an immune checkpoint that inhibits the excessive response of antigen-activated T cells to prevent autoimmune tissue damage. In chronic infections or cancers, lasting antigen exposure leads to permanent PDCD1 expression that can limit immune-mediated clearance of pathogens or degenerated cells. Consequently, blocking PDCD1 can enhance T cell function, which is the basis of cancer immune checkpoint therapy. We found that PDCD1 expression can increase within hours of temperature elevation in human leukemic and lymphoblastoid cell lines (such as Jurkat, THP1, HL-60, GM07062, and NK-92) and mouse lymphoid organs (e.g., thymus, spleen, and lymph nodes). Transcriptional upregulation of the *PDCD1* gene was associated with the binding of heat shock factor 1 (HSF1) to the promoter, and HSF1 knockout in HL-60 cells resulted in reduced *PDCD1* activation. Furthermore, a heat shock-dependent increase in glycosylated (and therefore active) PDCD1 protein levels was associated with PDCD1 exposure on the cell membrane and a reduction in the cytotoxic properties of NK-92 cells. Our observations suggest that the immune response could be attenuated in various physiological conditions accompanied by increased temperatures (infection, heat stroke, etc.). This observation may have clinical implications, and therefore, further research is warranted to understand the importance of fever and PDCD1 in various disease states, as well as their interaction with treatment.

## Introduction

The immune system has evolved to protect the organism from diseases. It must detect and react to pathogens, cancer cells, and foreign bodies, distinguishing them from healthy tissue in the body. During the immune response to defend against foreign invaders, innate (nonspecific) and adaptive (acquired) systems are activated. Innate immune cells are the ‘first responders’, arriving within hours to destroy pathogens through phagocytic or cytotoxic activities. These activities limit infection until a peak adaptive immune response is generated, normally around a week later. When eliminating a threat, the immune system must do so in a way that protects healthy cells and maintains self-tolerance. This action must be precisely controlled to ensure an adequate response (dysfunction of the immune system can cause autoimmune or inflammatory diseases and cancer), which is accomplished by repeatedly checking and balancing the immune responses. Among stimulatory and inhibitory checkpoint molecules, PDCD1 (programmed cell death protein 1, also known as PD1 or CD279) has received considerable attention for its role in maintaining T cell exhaustion, which is characterized by a gradual and progressive loss of T cell functions and can culminate in the physical deletion of the responding cells ^1^.

PDCD1 is a receptor found mainly on the surface of T cells (and to a lesser extent, on other blood and immune cells) ^2^. It is heavily glycosylated, which is critical for its biological functions ^3^. It can bind two ligands, PD-L1 (official symbol: CD274) or PD-L2 (PDCD1LG2), decreasing immune system reactivity and promoting self-tolerance (by suppressing pro-inflammatory activity) ^4^. PDCD1 signaling prevents excessive T cell activation during acute infections and maintains T cell exhaustion during chronic infections ^5^. PDCD1 can also negatively regulate B cell activation and proliferation ^6^. High and sustained expression of PDCD1 and its ligands is often observed during chronic infections and cancer ^7^. Tumor cells can exploit the PDCD1 signaling pathway to achieve immune evasion. It has been shown that blocking the PDCD1 signaling can improve T cell function and reduce viral load ^8^ and tumor burden ^9^. Consequently, recently developed inhibitors of the PDCD1 pathway have revolutionized cancer treatment for some patients. PDCD1 overexpression in T cells can also accompany other disease states, such as type 2 diabetes, which is associated with immune dysfunction and the development of cardiovascular disease ^10^. On the other hand, PDCD1 signaling is essential during pregnancy, and its blockade or deficiency is associated with embryo loss ^11^.

Several inflammatory reactions are induced in response to infection. In addition to the local inflammatory response, systemic defense reactions known as the acute phase reaction are triggered, which is characterized by pyrogenic fever, accelerated growth of peripheral leukocytes, circulating neutrophils, and their precursors. Fever can be defined as a controlled increase in body temperature as a result of an upward reset of the hypothalamic thermostat. This distinguishes fever from hyperthermia (sometimes called nonpyrogenic fever), e.g. heat stroke, which involves overheating of the body while the temperature set-point remains normal. In addition to the most common infectious etiology, fever can have a noninfectious cause (in which case it is usually chronic or recurrent). It can occur as a result of hypersensitivity reactions, autoimmune diseases, or malignancy ^12^. One of the benefits commonly attributed to pyrogenic fever is the enhancement of immunoprotective mechanisms (both innate and adaptive) during infection. Many conditions considered non-pyrogenic can also stimulate an inflammatory response. However, temperatures in the fever range can also have detrimental effects. Consequently, uncontrolled fever is associated with worse outcomes in patients with sepsis or neurological injuries ^13,14^.

Elevated body temperatures (regardless of the cause) lead to the activation of HSF1 (heat shock factor 1) ^15^, which is a major mediator of transcriptional responses to proteotoxic stress (including heat stress), frequently overexpressed in cancer ^16^. HSF1 is not only the central regulator of the heat shock response resulting in HSPs (heat shock proteins) synthesis and cytoprotection ^17^ but also plays a role in systemic thermoregulation ^18^, gametogenesis, development and other physiological and biological processes ^19,20^, including the replication cycle of many viruses ^21^. Furthermore, it regulates the expression of pivotal cytokines and early response genes ^22,23^. Intracellular HSPs serve as chaperones that can suppress apoptosis and inflammatory signaling. On the other hand, extracellular HSPs can provide the danger signals to enhance inflammation ^13,24,25^. Interestingly, in some cells, HSF1 can inhibit excessive synthesis of inflammatory cytokines, which, together with the upregulation of chaperone proteins, can prevent inflammation ^23^. In heat-sensitive cells, HSF1 can play the opposite role, that is, it induces apoptosis through upregulation of *PMAIP1* ^26^. Here, we found that HSF1 may also play a role in transcriptional upregulation of the *PDCD1* gene. Moreover, proteotoxic stress leads to the accumulation of glycosylated PDCD1 and its exposure on the cell surface, which may affect cellular functions.

## Materials and methods

### Cell lines

Human HL-60 (acute promyelocytic leukemia, promyeloblast) (RRID:CVCL_0002), Jurkat (acute T cell leukemia, T lymphoblast) (RRID:CVCL_0065), THP1 (acute monocytic leukemia, monocyte) (RRID:CVCL_0006), GM07062 (B-lymphoblastoid cell line from B lymphocyte) (RRID:CVCL_F143), NK-92 (natural killer) (RRID:CVCL_2142), U2OS (osteosarcoma) (RRID:CVCL_0042), RKO (colon carcinoma) (RRID:CVCL_0504), K562 (chronic myelogenous leukemia) (RRID:CVCL_0004), Raji (Burkitt’s lymphoma, B lymphocyte) (RRID:CVCL_0511), BJ1-hTERT (normal fibroblasts) (RRID:CVCL_6573), HEK293T (embryonic kidney, a line with neuronal cell characteristics) (RRID:CVCL_0063), HUVEC-C (umbilical vein endothelial) (RRID:CVCL_2959), RPMI-1788/CCL156 (B lymphocyte) (RRID:CVCL_2710), breast adenocarcinomas: MCF7 (RRID:CVCL_0031), T47D (RRID:CVCL_0553), CAL120 (RRID:CVCL_1104), MDAMB231 (RRID:CVCL_0062) cell lines were cultured in recommended media supplemented with 10% fetal bovine serum (FBS) (EURx, Gdansk, Poland). NK-92 medium was supplemented with 20 ng/ml of human recombinant IL-2 (Proteintech Group, Inc, Rosemont, IL, USA). Cells were routinely tested for mycoplasma contamination.

### Heat shock treatments

For heat shock, logarithmically growing cells or isolated PBMCs were placed in a water bath at the indicated temperature (38-43 °C) and for the specified time and allowed to recover for the indicated time in a CO_2_ incubator at 37 °C. The growth media were not replaced either before or after treatments. Adult (10–16-week-old), inbred FVB/N male and female mice were used for whole-body heat treatment (in a 42 °C water bath for 30 min) as previously described ^27^. Animal experiments were carried out according to Polish legislation and were approved by the Local Committee of Ethics and Animal Experimentation at the Medical University of Silesia in Katowice, Poland (Decision No. 129/2014 made on 17 December 2014) and by the Institutional Animal Care Policy of the Maria Skłodowska-Curie National Research Institute of Oncology (Gliwice, Poland).

### Functional HSF1 knockout using the CRISPR/Cas9 editing system

To remove the human HSF1 gene, we used a DNA-free system: Edit-R predesigned synthetic Human HSF1 sgRNAs (GGTGTCCGGGTCGCTCACGA in exon 1 on the minus strand, GTGGTCCACATCGAGCAGGG in exon 3 on the plus strand, and TCTCCCAGCTCAAACAGCAC in exon 13 on the minus strand), (Dharmacon/Horizon Discovery Ltd., Cambridge, United Kingdom), and eSpCas9 protein (Cat# ESPCAS9PRO, Merck KGaA) were introduced into HL-60 cells by electroporation using 4D-Nucleofector (program EO-100) and P3 Primary Cell 4D-Nucleofector™ X Kit (Cat#: V4XP-3032; Lonza, Basel, Switzerland) according to the protocol (the procedure was repeated twice with an interval of ∼0.5 minute). Single clones were obtained by limiting dilution on a 96-well plate. The efficiency of the HSF1 knockout was monitored by western blot and confirmed by sequencing (Genomed, Warszawa, Poland). In clone #16, 32 bp in exon 13 were deleted, and potentially a new stop codon located downstream could be used. Theoretically, a protein of 606 aa (instead of the wild-type 529 aa) can be produced, of which 484 aa are identical to the wild-type protein (only the C-terminal domain of HSF1 is affected). In clone #215, changes occurred between exon 1 and 3 (it was not possible to read the sequence accurately because the sequences from the two alleles of the gene were different).

### RNA isolation, cDNA synthesis, and RT-qPCR

Total RNA was isolated using the Universal RNA Purification Kit (EURx, Gdansk, Poland), digested with DNase I (Worthington Biochemical Corporation, Lakewood, NJ, USA), and cleaned with Clean-Up RNA Concentrator (A&A Biotechnology, Gdansk, Poland). RNA (1 μg) was converted into cDNA as described ^28^. Quantitative PCR was performed using a BioRad C1000 TouchTM thermocycler connected to the CFX-96 head (Bio-Rad Laboratories, Inc, Hercules, CA, USA). Each reaction was performed in triplicates using PCR Master Mix SYBRGreen (A&A Biotechnology, Gdansk, Poland). Expression levels were normalized against *HPRT1, GAPDH, ACTB*, and *HNRNPK*, or *TMEM43* in the case of human samples, and *Gapdh* and *Hnrnpk* in the case of mouse samples. The set of delta-Cq replicates (Cq values for each sample normalized against the geometric mean of reference genes) for control and tested samples were used for statistical tests and estimation of the p-value. Shown are the median, maximum, and minimum values of a fold change versus the untreated control. The primers used in these assays are described in Table S1.

### Chromatin Immunoprecipitation (ChIP) and ChIP-Qpcr

ChIP assay was performed according to the protocol of the iDeal ChIP-seq Kit for Transcription Factors (Diagenode, Denville, NJ, USA) using an anti-HSF1 antibody (Cat# ADI-SPA-901, RRID:AB_10616511, Enzo Life Sciences, Farmingdale, NY, USA), and the results were analyzed by qPCR as described in detail in ^29^. The sequences of the primers used are presented in Table S2.

### Protein extraction and Western blotting

Whole-cell extracts were prepared using RIPA buffer supplemented with Complete™ protease inhibitor cocktail (Roche, Indianapolis, IN, USA) and phosphatase inhibitors PhosStop™ (Roche). Extracts from adherent cells were prepared without trypsinization (directly on the culture dish). Proteins (20–30 μg) were separated on 10% SDS-PAGE gels and blotted onto a 0.45 μm pore nitrocellulose filter (GE Healthcare) using the Trans-Blot Turbo system (Thermo Scientific™ Pierce™ G2 Fast Blotter) for 10 min. Following primary antibodies were used: against PDCD1 (1:2,000 – 1:6,000; Cat# 66220-1-Ig, RRID:AB_2881611), PD-L1/CD274 (1:1,000; Cat# 82719-15-RR, RRID:AB_3086518), GAPDH (1:7,000, Cat#: 60004-1-Ig, RRID:AB_2107436), all from Proteintech (Rosemont, IL, USA), HSF1 (1:2,000; Cat# ADI-SPA-901, RRID:AB_10616511) and HSPA1A/HSP70 (1:5,000, Cat# ADI-SPA-810, RRID:AB_10616513), both from Enzo Life Sciences, ACTB (1:25,000, Cat# A3854, RRID:AB_262011, Merck KGaA). When necessary, the primary antibody was detected by an appropriate secondary antibody conjugated to horseradish peroxidase (Thermo Fisher Scientific, Waltham, MA, USA) and visualized by an ECL kit (Thermo Fisher Scientific) or WesternBright Sirius kits (Advansta, Menlo Park, CA, USA). Imaging was performed on X-ray film or a G:BOX chemiluminescence imaging system (Syngene, Frederick, MD). The blots were subjected to densitometric analyses using Image Studio Lite v. 5.2.5 software to calculate relative protein expression after normalization with loading controls (statistical significance of differences was calculated using a T-test).

### Measurement of PDCD1 expression on the cell surface by flow cytometry

One million cells per sample were used, and all steps were performed in a volume of 50 µl and at room temperature. Cells were washed twice with PBS and centrifuged for 2 minutes at 2,000 rpm. The cell pellet was resuspended in PBS, and CoraLite® Plus 488-conjugated antibodies were added: 0.4 µg anti-PDCD1 (Proteintech, Cat# CL488-66220; RRID:AB_2883287) or 0.4 µg mouse IgG1 isotype control (Proteintech, Cat# CL488-66360-1; RRID:AB_2934458). Cells were incubated for 30 minutes in the dark, then washed twice with PBS, resuspended in PBS, and analyzed by flow cytometry (in parallel with unstained cells).

### NK-92 cytotoxicity

NK-92 cells were left untreated or subjected to heat shock for 1 hour at 42 °C or 42.5 °C. After ∼20 h, cell viability was assessed using trypan blue solution. 200,000 viable NK-92 cells were mixed in a 1:1 ratio with viable target cells (in a 2 ml medium dedicated to NK-92 cells) and seeded in a well of a 6-well plate. As a control, only 200,000 viable target cells were seeded in an NK-92 cell medium. After 24 hours of culture, the medium was removed, the wells were washed with PBS, and the remaining adherent cells were fixed with frozen methanol and dried. The cells were stained with crystal violet, rinsed with distilled water, and dried again. The bound crystal violet was extracted using 1 ml of 10% acetic acid, and absorbance was measured at 595 nm. In three biological replicates, two technical replicates each were performed.

### Statistical analyzes

For each dataset, the normality of the distribution was assessed using the Shapiro-Wilk test. Depending on data distribution, the homogeneity of variances was verified by the Levene test or Brown-Forsythe test. Outliers were determined using the Tuckey criterion and the QQ plot. For analysis of differences between compared groups with normal distribution, the quality of the mean values was verified by the ANOVA test with a pairwise comparison done with the HSD Tukey test or Games-Howell test depending on the homogeneity of variance. In the case of non-Gaussian distribution, the Kruskal–Wallis ANOVA was applied to verify the hypothesis on the equality of the medians with the Conover-Iman test or Dunn test for pairwise comparisons. P = 0.05 was selected as a statistical significance threshold.

## Results

Analyzing available ChIP-seq data from MCF7 breast cancer cells (GSE137558 ^29,23^) and U2OS osteosarcoma cells (GSE60984 ^22^), we noticed that the temperature elevation led to HSF1 binding to the *PDCD1* gene promoter (in the heat shock element, HSE, located ∼670 bp upstream of the transcription start site; Fig. S1A). Additionally, RNA-seq data (ArrayExpress, acc. No. E-MTAB-13903) indicates that HSF1 binding to the *PDCD1* promoter may correlate with increased transcriptional activity of the gene in MCF7 cancer cells, but not in noncancerous MCF10A breast epithelial cells (Fig. S1B). This suggests that *PDCD1* expression may be up-regulated by HSF1 during the heat shock response and that even non-immune cells may respond to such stimulation. Therefore, we checked the activation of *PDCD1* transcription by heat shock in a panel of cell lines of different origins. Initial studies showed increased *PDCD1* expression in several leukemia cell lines: HL-60 (promyeloblasts from a patient with acute promyelocytic leukemia) K562 (hematopoietic cells from a patient with chronic myeloid leukemia at a blast crisis), Raji (B lymphocytes from a patient with Burkitt’s lymphoma), and Jurkat (T lymphoblasts from a patient with acute T cell leukemia), but also in HEK293T (a line with neuronal cell characteristics, derived from the embryonic kidney). No expression was found in BJ1-hTERT (normal fibroblasts) and HUVEC-C (umbilical vein endothelial cells) (Fig. S1C). Further more detailed studies confirmed the increase in *PDCD1* transcript levels after heat shock in some cells of hematopoietic origin: HL-60, Jurkat, THP1 (monocytes from a patient with acute monocytic leukemia), GM07062 (B-lymphoblastoid), and NK-92 (natural killers from a patient with malignant non-Hodgkins lymphoma) cells (Fig. 1A, F). However, no meaningful heat shock-induced *PDCD1* upregulation was observed in U2OS (osteosarcoma) and RKO (colon cancer) cells (Fig. 1A).

**Fig. 1.**
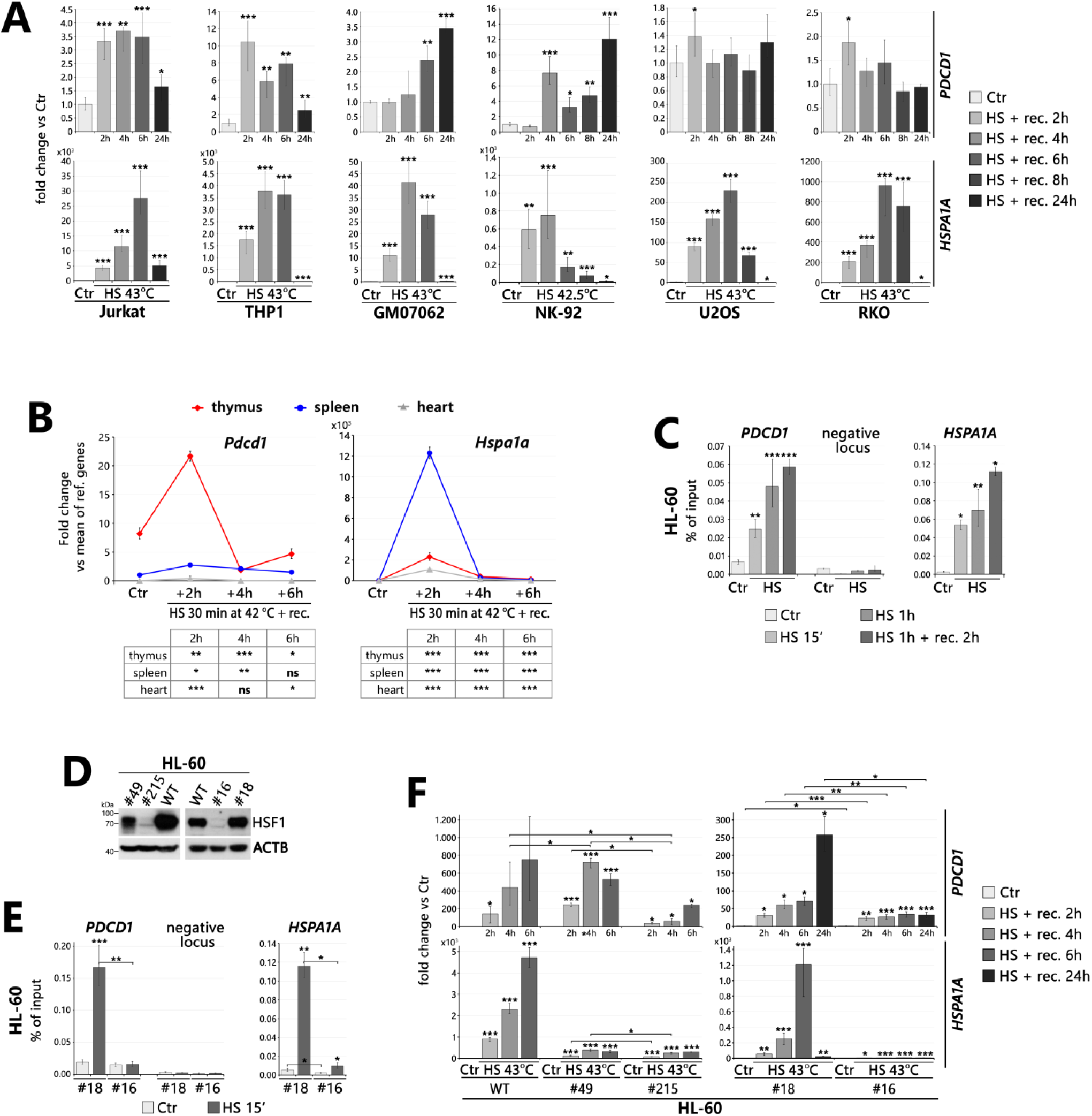
Up-regulation of *PDCD1* transcription after heat shock in human cell lines and mouse tissues can be mediated by HSF1. (**A**) *PDCD1* and *HSPA1A* (positive control for the heat shock response) transcript levels after heat shock (HS) were analyzed by RT-qPCR in human cell lines. (**B**) *Pdcd1* and *Hspa1* expression after heat shock (HS *in vivo*) was analyzed by RT-qPCR in the mouse thymus, spleen, and heart. The readings were normalized against the mean of reference genes and presented versus Ctr in the spleen. Statistical significance versus corresponding Ctr is shown in the tables below. (**C**) HSF1 binding to the *PDCD1* promoter was analyzed by ChIP-qPCR in the HL-60 cell line, untreated (Ctr) and after heat shock (HS at 43 °C). HSF1 binding to the negative locus and *HSPA1A* promoter were shown as negative and positive controls, respectively. (**D**) Western blot analysis of HSF1 levels in HL-60 cells: wild type (WT) and individual clones obtained after CRISPR/Cas9 editing (with HSF1 deficiency: #49, #215, #16, and unaltered: #18). (**E**) HSF1 binding to the *PDCD1* promoter (analyzed by ChIP-qPCR) confirming the absence of heat-induced (HS at 43 °C) binding in the HSF1-deficient clone (#16). HSF1 binding to the negative locus and *HSPA1A* promoter were shown as negative and positive controls, respectively. (**F**) RT-qPCR analysis of *PDCD1* and *HSPA1A* levels in WT and modified HL-60 cells. *** p < 0.0001, ** p < 0.001, * p < 0.05 (significance of differences).

The mouse *Pdcd1* gene promoter also contains HSE (Fig. S1A). Therefore, we checked the gene activation after heat shock in the mouse thymus (where T cells mature) and spleen (which acts primarily as a blood filter). *Pdcd1* is expressed mainly in T cells, therefore, its expression was higher in the untreated thymus than in the untreated spleen and was undetectable in the heart (Fig. 1B). Nevertheless, in both the thymus and spleen (and even slightly in the heart), *Pdcd1* was upregulated after heat shock. We postulate that heat shock-induced upregulation of *PDCD1* transcription may be mediated by HSF1 since HSF1 binding to the *PDCD1* promoter was increased under such conditions, as demonstrated by ChIP-qPCR in HL-60 cells (Fig. 1C, E). Moreover, *PDCD1* transcription in HSF1 knockout cell lines was not induced after heat shock as efficiently as in HSF1-containing lines. (Fig. 1D – F).

Heat shock also induced PDCD1 protein accumulation. PDCD1 can be detected in two forms: nonglycosylated and glycosylated. Glycosylation was shown to be critical for maintaining the stability of the PDCD1 protein, cell surface localization, and mediating its interaction with PD-L1 ^3^, therefore, it is considered to be the active form. The antibody we used (see Fig. S2A and B for its specificity) detected one or both forms of the protein in different human cells and mouse thymus (Fig. S2C). The increase in PDCD1 levels was observed after heat shock in human cell lines of hematopoietic origin: HL-60, Jurkat, GM07062, THP1, and NK-92 (Figs 2A and S2D), but also in the mice lymph nodes (Fig. 2A), spleen, and Peyer’s patches (not shown). Moreover, another proteotoxic agent, bortezomib (proteasome inhibitor), induced the accumulation of glycosylated PDCD1 (Fig. S2E). It is worth noting that the most effective treatments (heat shock at 43 °C, 32 nM bortezomib) led to higher mortality in some cell lines.

**Fig. 2.**
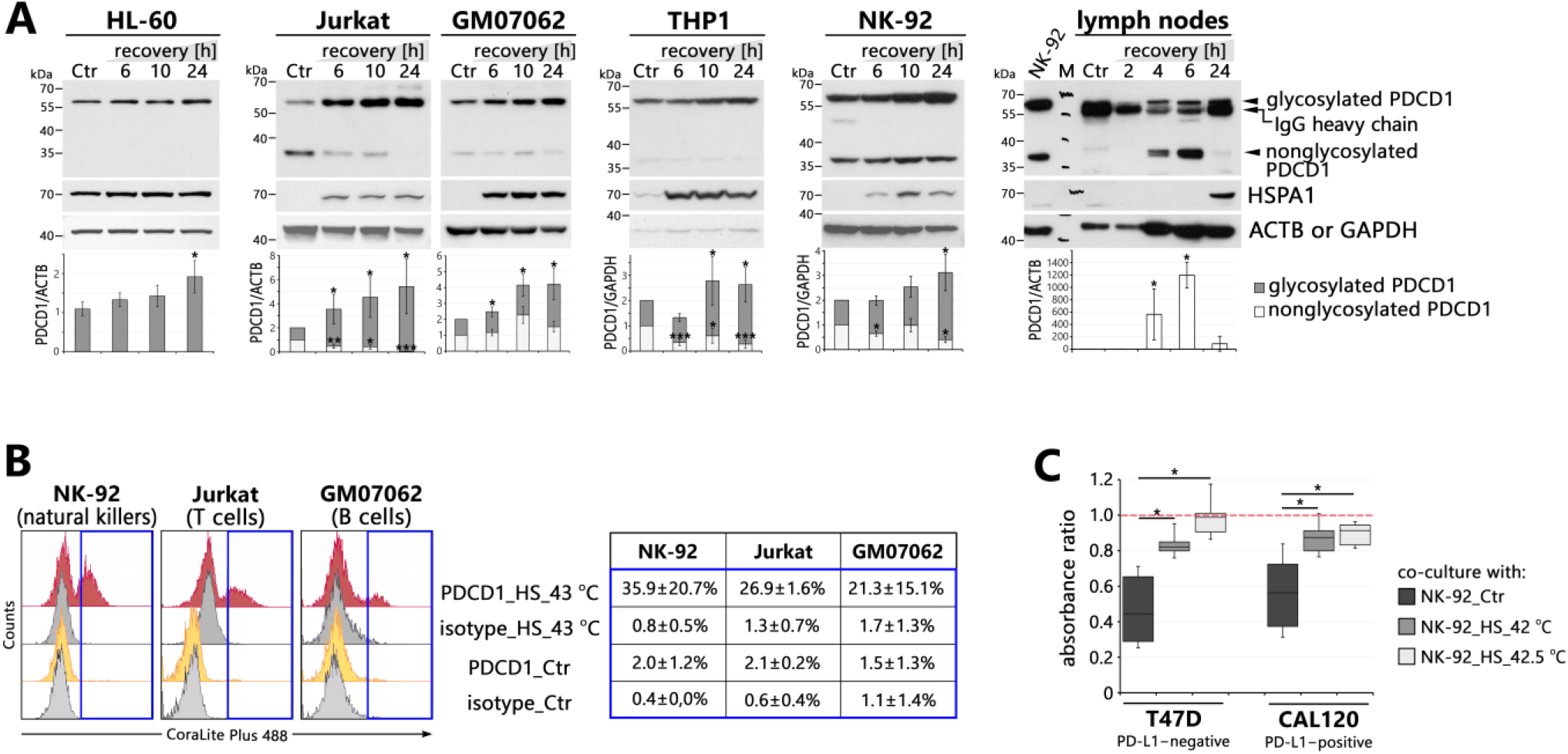
Heat shock treatment can increase PDCD1 protein levels, which may have functional consequences. (**A**) Western blot analyses in human cell lines and mouse lymph nodes heat-shocked (HS, 1 h at 43 °C; 42.5 °C in the case of NK-92 and 42 °C – mouse tissues) with the indicated recovery time. HSPA1 and ACTB or GAPDH were used as positive controls for the HS response and loading controls, respectively. Graphs show the results of densitometric analyses (in the case of mouse tissues, the glycosylated form was not analyzed due to the overlap of the IgG signal detected by the secondary antibody). (**B**) Cell surface staining of PDCD1 was analyzed by flow cytometry in untreated (Ctr) and heat-shocked (HS, 1h at 43 °C and 12h recovery) cells. The counts (y-axis not to scale) from the blue squares areas are shown (as % of parent) in the table on the right. (**C**) Cytotoxicity of untreated and heat-shocked NK-92 cells against T47D and CAL120 cells. The absorbance ratio: absorbance of the crystal violet-stained target cells after 24 h co-culture with NK-92 was normalized versus absorbance of unattacked cells, which is 1.0 (red dashed line). Boxplots represent the median, upper and lower quartiles, maximum, and minimum. *** p < 0.0001, ** p < 0.001, *p<0.05 (significance of differences).

Heat shock primarily increased the levels of glycosylated (and therefore active) PDCD1 protein in human cell lines, which was associated with its exposure on the cell membrane as shown in NK-92, Jurkat, and GM0702 cells (Fig. 2B). To investigate the functional consequences of heat shock, we first analyzed the cytotoxicity of NK-92 cells against the four breast cancer cell lines: MCF7, T47D (PDL1-negative), CAL120, and MDAMB231 (PDL1-positive) (Fig. S2F). Only T47D and CAL120 cells were attacked by NK-92 (Fig. S2G), but the cytotoxic properties of NK-92 were reduced after heat shock (Fig. 2C).

## Discussion

In addition to epigenetic mechanisms, the *PDCD1* expression is regulated by distinct transcription factors in different immune cells, and it is regulated by differential mechanisms during acute and chronic infections. It can be activated by NFATC1, RBPJ, STAT3, STAT4, STAT1 (ISGF3), FOXO1, FOS, and NF-κB and repressed by PRDM1 (BLIMP1) and TBX21 (T-bet) transcription factors in different circumstances ^30^. Here, we showed that in some cells, the *PDCD1* expression can also be upregulated at elevated temperatures by HSF1. The levels of the active (glycosylated) form of the PDCD1 protein increased, and the protein was exposed on the external side of the membrane a few to several hours after heat shock. This can have functional consequences during the body’s normal response to infections, but also in chronic inflammatory and autoimmune diseases, sepsis, pregnancy, cancer and anticancer therapy, organ transplantation, etc., if accompanied by severe fever.

Typically, 42/43 °C is applied to activate HSF1 in experimental studies. However, HSF1 activation and *HSP* gene expression have also been shown to occur at temperatures in the febrile range (typically 38.5–41 °C) in experimental and clinical studies ^15^. Interestingly, some cell types and tissues activate HSF1 at lower temperatures than others. For example, in T cells (where PDCD1 signaling is best studied), HSF1 is activated as early as 39 °C ^31^. It was shown that HSF1 plays a role in facilitating the proliferation after antigen-mediated activation of T cells, B cells, and hematopoietic stem cells at fever temperatures ^32,33^. In addition, the proliferation of antigen-activated T cells was also HSF1-dependent at non-febrile temperatures (but only in the spleen, not in the lymph node microenvironment) ^33^. Our results suggest that HSF1 may also exert another (possibly opposite) effect on immune cells by increasing the expression of PDCD1.

In addition to increased levels of *Pdcd1*/PDCD1 at elevated temperatures in the mouse lymphoid organs (suggesting activation in T cells), we found its increased levels also in cultured cell lines of hematopoietic origin. PDCD1 expression is known to be induced in T cells and B cells after their activation through antigen receptors ^34^, that is, a few days after infection, and is thought to attenuate the immune response ^4,35^. Fever is usually the first sign of infection. Thus, the upregulation of PDCD1 due to elevated temperatures accompanying infection is more likely to occur in T and B cells before their antigen-mediated activation (thus, in naive cells). It was already shown that PDCD1 can be transiently expressed in naive, virus-specific CD8 T cells shortly after acute virus infection, but this upregulation was driven predominantly by antigen receptor signaling. During the naive-to-effector CD8 T cell transition, PDCD1 also had an inhibitory role (and its blockade enhanced effector function and resulted in faster clearance of infection) ^36^. Our finding that elevated temperatures (even without infection) upregulate PDCD1 expression points to a new mechanism of decreasing immune system reactivity and promoting self-tolerance already before the development of an acquired immune response. This also means that fever may not only increase the immune system’s effectiveness during infection ^13^ but simultaneously may be involved in quenching (or balancing) the immune response. Alternatively, the action of PDCD1 may be different upon such activation, but its functional consequences require further studies.

There are few reports on the effects of fever on T cells. They showed a rather stimulating effect and have not been linked to PDCD1 expression. It was shown that temporary exposure of naive CD8+ T cells to elevated temperatures (39.5 °C) before antigen exposure resulted in a greater percentage of cells, which subsequently differentiated into effector cells ^37^. Also, fever-range temperatures modulated naive CD4 T cell differentiation (shifting the Th1/Th2 balance toward the Th2 phenotype, which is known to enhance the B cell-mediated antibody responses), while antigen-presenting cell (APC)-mediated CD4 T cell activation at elevated temperatures did not lead to Th2 differentiation ^38^. Furthermore, fever promoted T cell trafficking to lymphoid organs and inflamed tissues, which was beneficial and enhanced immune surveillance during infection ^39^. Studies have also demonstrated a pathogenic mechanism in which fever promotes autoimmune diseases by regulating the differentiation and pathogenicity of Th17 cells ^40^.

Although PDCD1 is expressed mainly in T cells, some reports indicate that it may also act as an immune checkpoint for innate lymphoid populations (e.g. NK cells, myeloid cells, monocytes and macrophages, etc.). PDCD1 has been shown to undergo a consistent basal expression on all circulating human NK cells and its blockade enhanced NK cell natural cytotoxicity ^41^. In addition, heat shock (42 °C) transiently inhibited human NK cells’ cytotoxicity ^42,43^, which is in line with our results. On the other hand, hyperthermia below 40 °C has been shown to have stimulatory effects on NK cells ^43^. These data suggest that hyperthermia can have immunosuppressive or stimulatory effects on NK cells, depending on the treatment protocol. Interestingly, PDCD1 is expressed in some tumor cells and our results suggest that this can be mediated by proteotoxic stress. PDCD1 has been shown to promote tumorigenesis in melanoma, hepatic carcinoma cells, pancreatic ductal adenocarcinoma, thyroid cancer, glioblastoma, and triple-negative breast cancer ^44,45^, while in lung cancer and colon cancer cells, it appears to play a different role, as tumor cell proliferation is induced when PDCD1 is blocked ^46,47^. This intrinsic PDCD1 expression in cancer cells may impact the clinical outcome of immunotherapy. Such PDCD1 acts independently of adaptive immunity and its action may explain the different therapeutic effects of anti-PDCD1 treatment and provide critical information for use in combined anti-tumor approaches.

Fever can also be one of the side effects of the anti-cancer therapy. On the other hand, hyperthermia is used to target cancer cells and their surrounding environment and has the potential for cancer therapy in conjunction with other treatments ^48,49,25,50^. Therefore, it would be appropriate to investigate the effect of fever/hyperthermia and transient PDCD1 expression (both on immune and cancer cells) on the efficacy of anticancer treatment, in particular immunotherapy.

To sum up, our results show for the first time that the expression of PDCD1 is positively regulated by elevated temperatures in various cell lines derived from hematological malignancies and murine immune organs such as the thymus, spleen, and lymph nodes. Heat shock-induced expression of PDCD1 was observed at both mRNA and protein levels. In the latter case, the accumulation of nonglycosylated and/or glycosylated (active) forms of the protein was observed, depending on the cell line/tissue and conditions of heat shock. Our results suggest that fever (via PDCD1 accumulation) may be involved in the attenuation of the immune response in various physiological states accompanied by increased temperatures (infection, heat stroke, etc.). For this reason, fever (as well as pharmacological fever reduction) can have unexpected consequences depending on the medical condition. This observation may have clinical implications and therefore, further research is warranted to understand the importance of fever on immune function in various disease states as well as its interaction with treatment.

## Supporting information

Original Images for Blots

## Conflict of Interest

The authors declare that the research was conducted in the absence of any commercial or financial relationships that could be construed as a potential conflict of interest.

## Author Contributions

Conceptualization – AT-J, PJ, WW; Data curation – PJ; Formal analysis – AT-J, PJ, WW; Funding acquisition – AT-J, WW; Investigation – AT-J, PJ, KM, KS, NV, MB, JM, WW; Methodology – AT-J, PJ, KM, KS, NV, WW; Project administration – AT-J, WW; Resources – WF, WW; Supervision – PJ, MO, WW; Validation – AT-J, PJ, WF; Visualization – KM, MO, WW; Writing – original draft – AT-J, WW; Writing – review & editing – MO, WW. All authors contributed to the manuscript revision, read, and approved the submitted version.

## Funding

This research was funded by NIO-PIB, internal grant SN/MGW11/2024 to AT-J and National Science Centre, Poland, grant 2021/43/B/NZ3/02161 to WW. For the purpose of Open Access, the author has applied a CC-BY public copyright license to any Author Accepted Manuscript (AAM) version arising from this submission.

## Acknowledgments

We thank Marek Rusin and Ryszard Smolarczyk for providing the protocols for culturing the NK-92, THP1, GM07062, and HUVEC-C cell lines. A preprint of the manuscript is available in the bioRxiv repository (https://doi.org/10.1101/2025.05.22.652424).

## Notes

### Competing Interest Statement

The authors have declared no competing interest.

### Summary of Updates

Minor changes to the summary and text. Additional supplementary file with original images for blots.

