## Supplementary Material for "PDCD1 expression increases at elevated temperatures"

### Supplementary materials and methods

**PDCD1 cloning and overexpression.** Human *PDCD1* coding sequence was amplified (fragment 48 – 930 nt from NM_005018.3) by PCR on cDNA from peripheral blood mononuclear cells as template and cloned into pLVX-puro vector (Cat. No. 632164; Takara/Clontech, Saint-Germain-en-Laye, France) between *EcoRI* and *ApaI* sites using NEBuilder HiFi DNA Assembly Master Mix (New England Biolabs, Ipswich, MA, USA). The DNA sequence of the recombinant plasmid was confirmed by sequencing (Genomed, Warszawa, Poland). The pLVX-puro (empty) and pLVX-PDCD1 vectors were introduced into HEK293T packaging cells using Lenti-X™ Packaging Single Shots (VSV-G) (Takara, Cat. No. 631275), and the viruses were produced and concentrated (Lenti-X™ Concentrator; Takara, Cat. No. 631231). MCF7 cells were transduced in the presence of polybrene (8 μg/ml) while Jurkat cells – using RetroNectin® (Takara, Cat. No. T100A) coated plates (10 μg/cm²). Puromycin was used to eliminate unmodified cells (1 μg/ml for one week in the case of MCF7 cells, 0.5 μg/ml for two weeks in the case of Jurkat cells). All procedures were performed according to the user manuals.

**Phytohemagglutinin, bortezomib, and IFNγ treatments.** Stock solution (1.6 mM) of bortezomib (Cat# S1013, Selleckchem, Houston, TX, USA) was prepared in DMSO. Working solutions (8, 16, 32 nM) were prepared fresh before each experiment in a culture medium (without antibiotics). Control cells were incubated with a medium containing DMSO. Cells were treated for 24 hours. Phytohemagglutinin (PHA-M, Cat# L2646, Merck KGaA, Darmstadt, Germany) was prepared in sterile water (25 mg/ml). Working solutions (10, 100, 1.000 ng/ml) were prepared fresh before each experiment in a culture medium. Interferon gamma (IFNγ, Cat# 130-096-872, Miltenyi Biotec B.V. & Co. KG, Germany) was prepared in sterile water (100ng/ml). Working solutions (1ng/ml) was prepared fresh before each experiment in a culture medium.

### Supplementary Figures

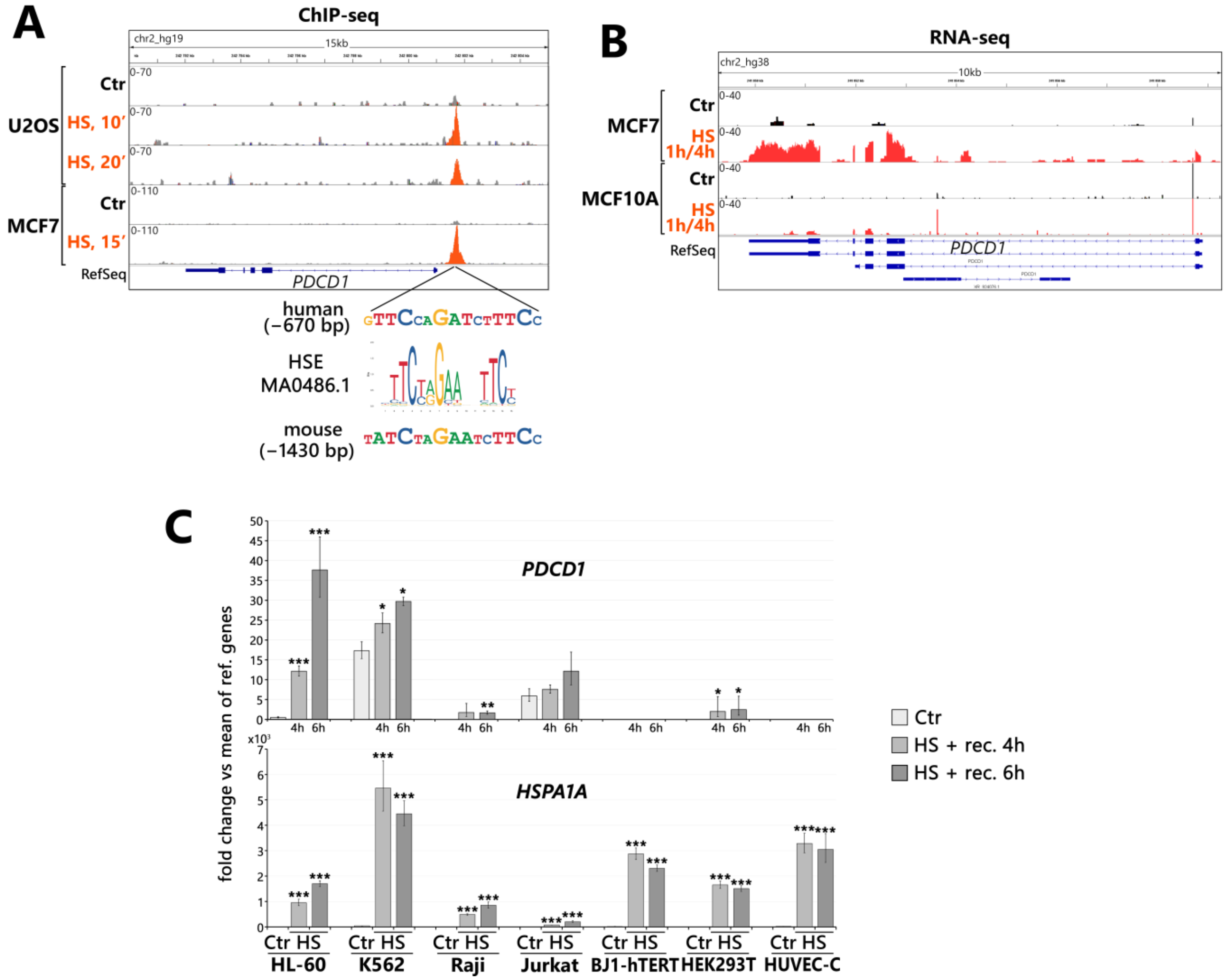


**Fig. S1.** Heat shock treatment causes increased binding of HSF1 to the *PDCD1* promoter and may result in transcriptional activation. (**A**) HSF1 binding to the *PDCD1* promoter in untreated (Ctr) and heat-shocked (HS at 43 °C) U2OS (osteosarcoma) and MCF7 (breast adenocarcinoma) identified by ChIP-seq (GSE60984, GSE137558: GSM4081759 and GSM4081762) (peaks are shown in the integrative genomic viewer, IGV). The heat shock element (HSE) sequence logo from the JASPAR database (MA0486.1) is shown below the HSE sequence present at the HSF1 binding site, and above the HSE present in the mouse *Pdcd1* promoter. (**B**) RNA-seq results (coverage 200M reads/sample; ArrayExpress, acc. No. E-MTAB-13903) showing *PDCD1* gene expression in untreated (Ctr) and heat shocked (HS at 43 °C for 1h and 4h of recovery) MCF7 and MCF10A (non-tumorigenic mammary epithelial) cells (reads shown in IGV). The scale for each sample is shown in the left corner. (**C**) *PDCD1* expression after heat shock (HS for 1h at 43 °C), was analyzed by RT-qPCR in human cell lines of different origin. *HSPA1A* expression was shown as a positive control for the heat shock response. Due to the frequent lack of expression in untreated cells (Ctr), the readings were normalized against the mean of reference genes (*GAPDH*, *ACTB*, *HPRT1*, *HNRNPK*) and presented versus Ctr in HL-60 cells. *** $p < 0.0001$, ** $p < 0.001$, * $p < 0.05$ (significance of differences).

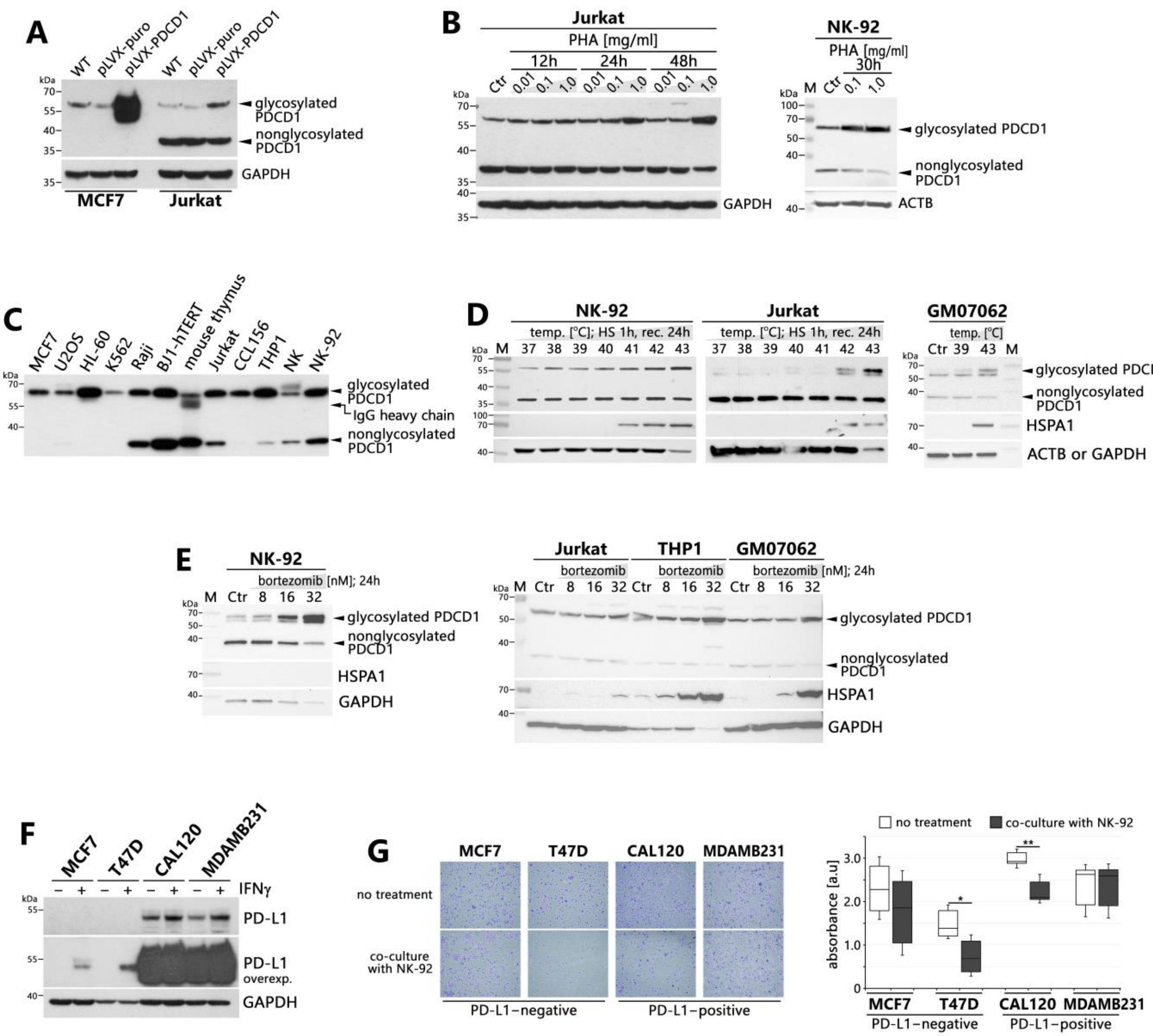


**Fig. S2.** Western blot analyses of PDCD1 and PD-L1 levels. Tests for specificity of the antibody: (**A**) overexpression of PDCD1 in MCF7 and Jurkat cells; (**B**) PDCD1 glycosylation stimulated with phytohemagglutinin (PHA) in Jurkat and NK-92 cells. (**C**) PDCD1 levels in a panel of human cell lines, isolated NK cells, and mouse thymus. (**D**) Temperature-dependent up-regulation of PDCD1 levels. Cells were heat shocked (HS) for 1 hour with indicated temperatures and recovered for 24h. (**E**) Effect of bortezomib (proteasome inhibitor) treatment on PDCD1 levels. Cells were treated for 24 hours with the indicated concentrations. HSPA1 and ACTB (or GAPDH) were used as positive controls for heat shock response and loading controls, respectively. (**F**) PD-L1 levels in breast cancer cell lines untreated and treated with IFNγ, which is known to upregulate PD-L1. (**G**) Tests of NK-92 cytotoxicity against selected breast cancer cell lines. Representative images of target cells stained with crystal violet are shown on the left, and a summary of cell survival (absorbance of crystal violet in arbitrary units, a.u., from three biological replicates) on the right. ** $p < 0.001$, * $p < 0.05$ (significance of differences).

### Supplementary Tables

**Table S1.** RT-qPCR primers for gene expression analyses.

| Gene symbol | RefSeq | forward primer sequence | reverse primer sequence |
|---|---|---|---|
| ***HSPA1A*** | NM_005345.5 | agctggagcaggtgtgtaaccc | aaaaacagcaatcttggaaaggccc |
| ***Hspa1a*** | NM_010479.2 | acaagagaagcagagcgagc | atcgccgtgttcttggccat |
| ***PDCD1*** | NM_005018.3 | ttccagtggcgagagaagacc | ggccaagagcagtgtccatc |
| ***Pdcd1*** | NM_008798.3 | ttgacacacggcgcaatgac | gccttgaaaccggccttctg |
| ***ACTB*** | NM_001101.5 | agagcctcgcctttgccgat | ttgcacatgccggagccgtt |
| ***GAPDH*** | NM_002046.7 | ttccatggcaccgtcaaggc | tgcaaatgagccccagccttct |
| ***Gapdh*** | NM_008084.3 | tggtgaagcaggcatctgagg | catgaggtccaccaccctgt |
| ***HNRNPK*** | NM_002140.4 | atgctgtcctcattccactgac | cgcgacggtcatcaaacatca |
| ***Hnrnpk*** | NM_001301341.1 | tggcatggttgggttcagtgc | ccacggccccctgcataagaat |
| ***HPRT1*** | NM_000194.3 | gccctggcgtcgtgattagt | tgatggcctcccatctcctt |
| ***TMEM43*** | NM_001407274.1 | cttgtggtgtctcccgacag | ttggtacatctccacgtgcc |

**Table S2.** ChIP-qPCR primers for HSF1 binding analyses.

| Gene symbol | RefSeq | forward primer sequence | reverse primer sequence |
|---|---|---|---|
| ***HSPA1A*** | NC_000006.12 | cggcactctggcctctgatt | gacccgccttttcccttctg |
| ***PDCD1*** | NC_000002.12 | gctaatcggctccccctcaaa | caggcatcacacggtggaaag |
| *negative control locus* | NC_000012.12 | tggcaccatgcttctttaagtcc | agtttgacaagttcaagcacc |
