## Supplementary material for "PDCD1 expression increases at elevated temperatures": Original Images for Blots

**Figure 1D**

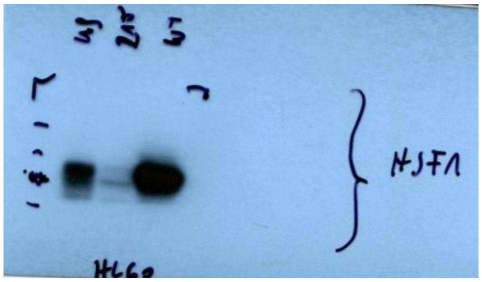

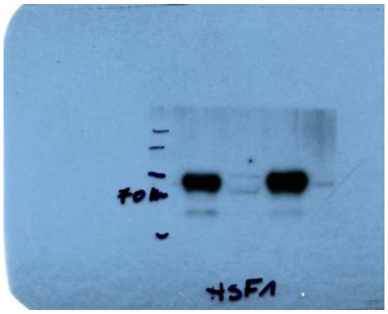

HSF1, HL-60

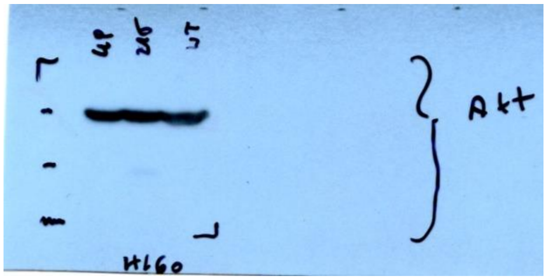


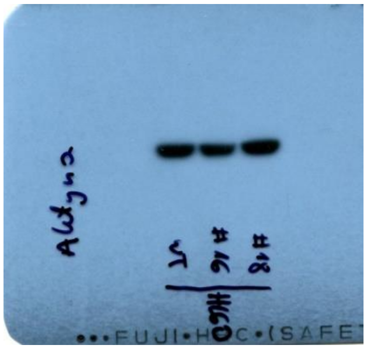

ACTB, HL-60

**Figure 2A**

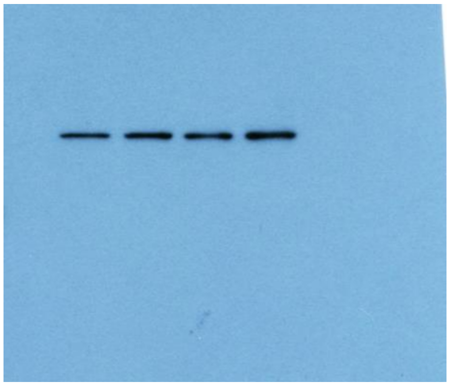

PDCD1, HL-60

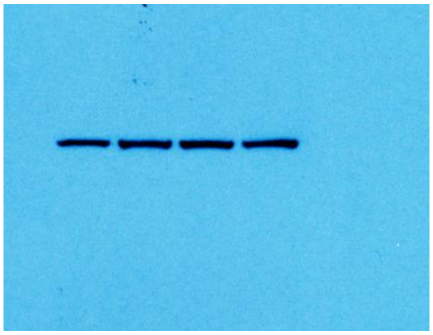

HSPA1, HL-60

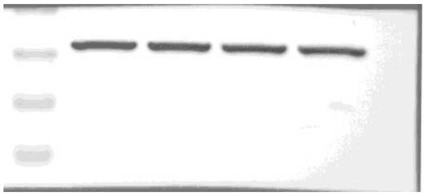

ACTB, HL-60

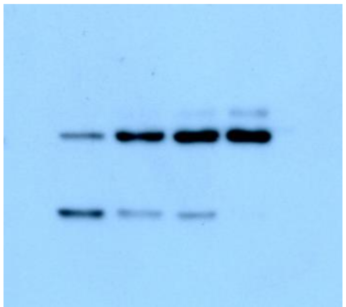

PDCD1, Jurkat

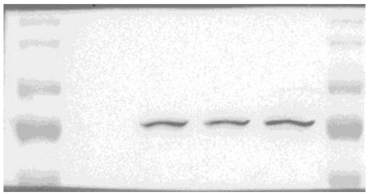

HSPA1, Jurkat

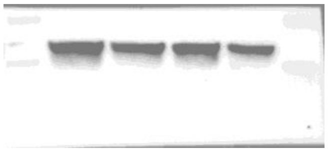

ACTB, Jurkat

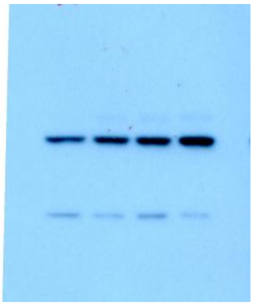

PDCD1, GM07062

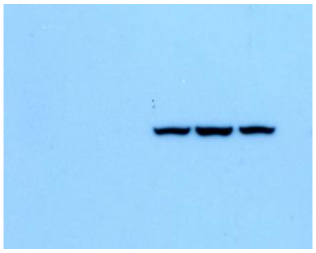

HSPA1, GM07062

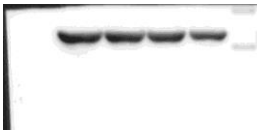

ACTB, GM07062

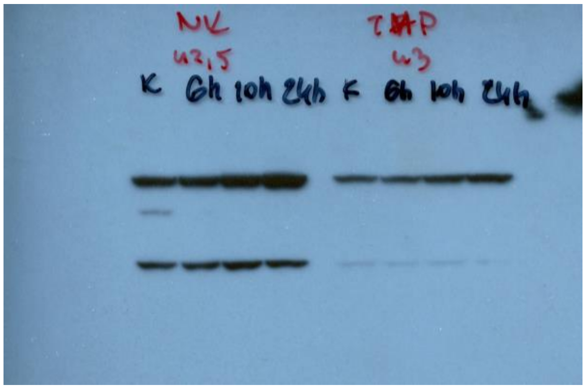


PDCD1, NK-92 and THP1

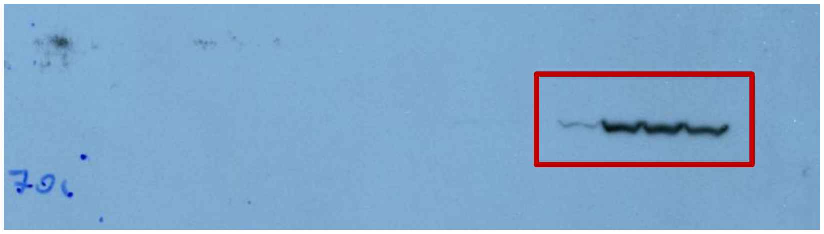

HSPA1, THP1

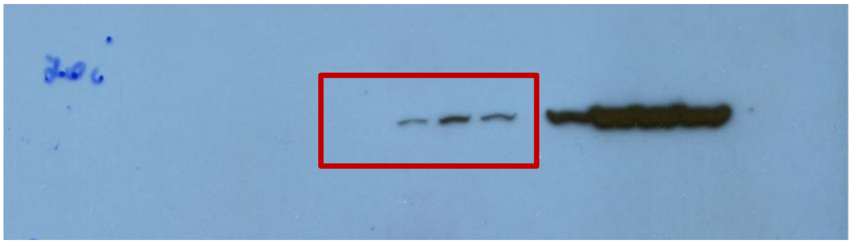

HSPA1, NK-92

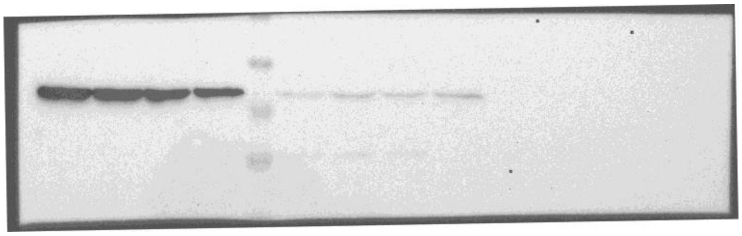

GAPDH, NK-92 and THP1

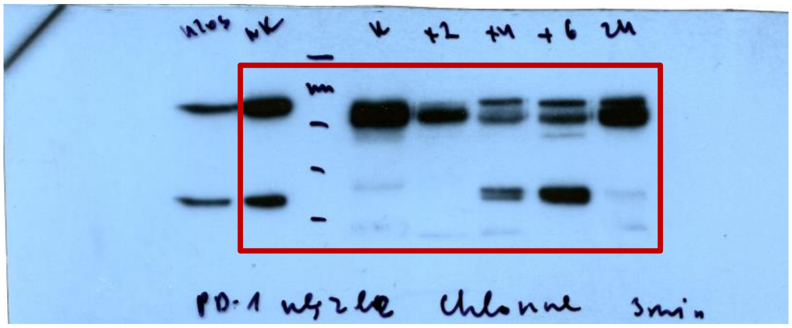

PDCD1, lymph nodes

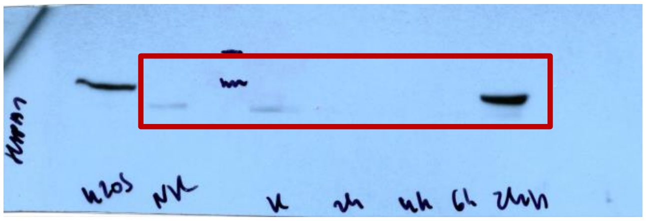

HSPA1, lymph nodes

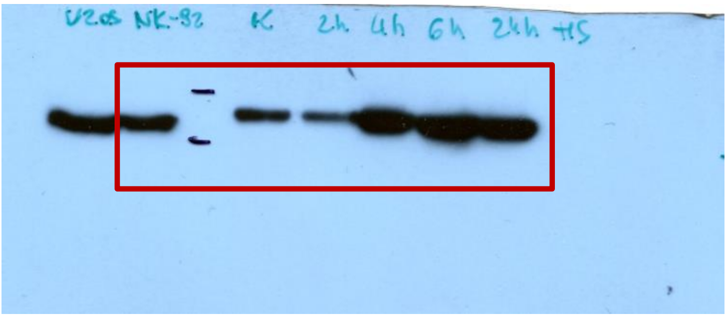


ACTB, lymph nodes

### Supplementary Figures

#### Figure S2A

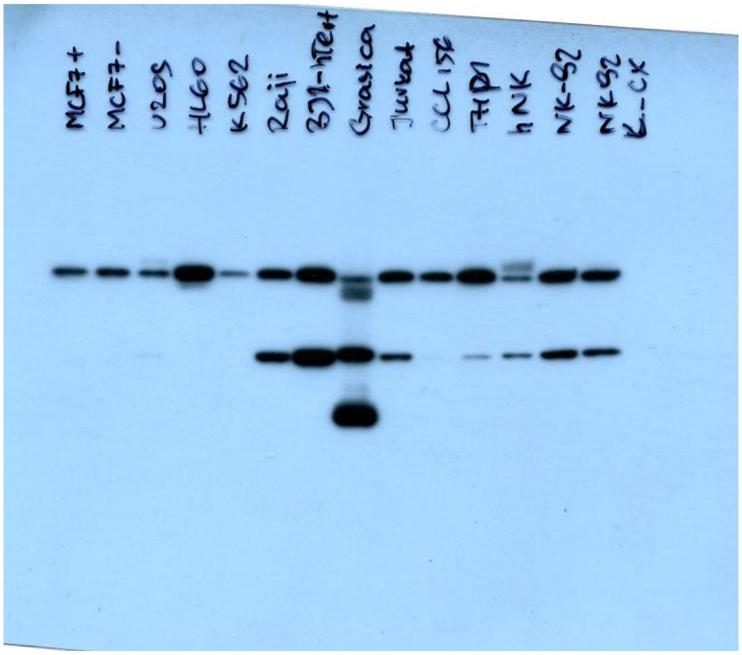


PDCD1

#### Figure S2B

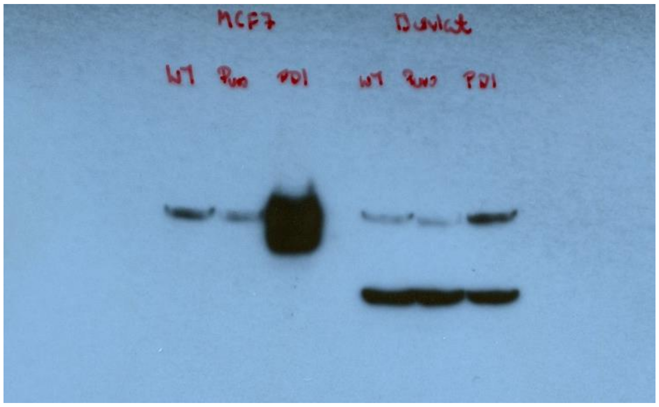


PDCD1, MCF7, Jurkat

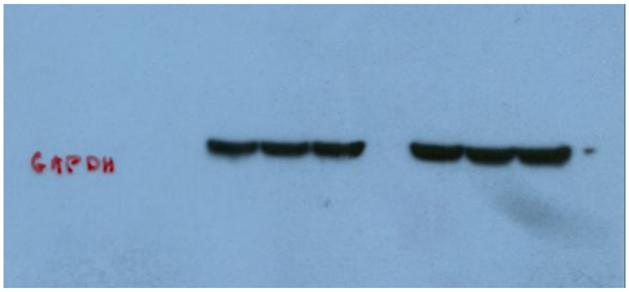


GAPDH, MCF7, Jurkat

**Figure S2C**

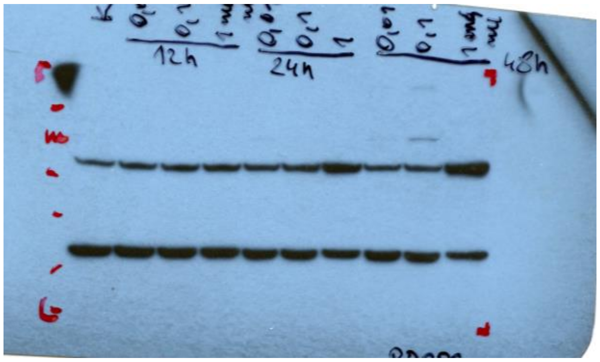


PDCD1, Jurkat

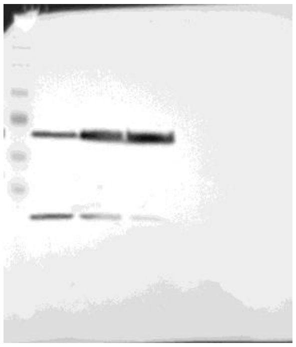

PDCD1, NK-92

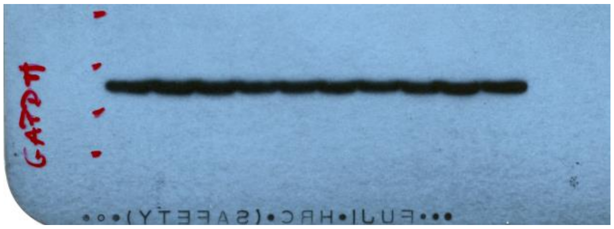


GAPDH, Jurkat

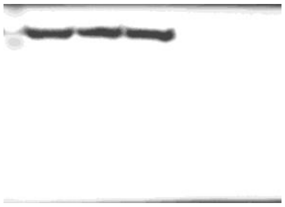

ACTB, NK-92

**Figure S2D**

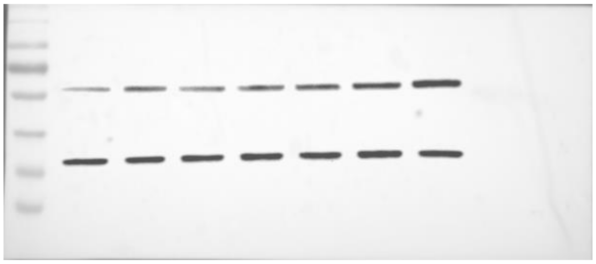

PDCD1, NK-92

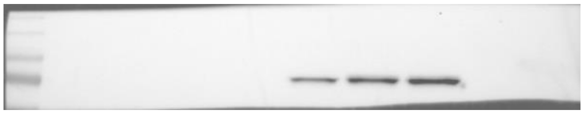

HSP70, NK-92

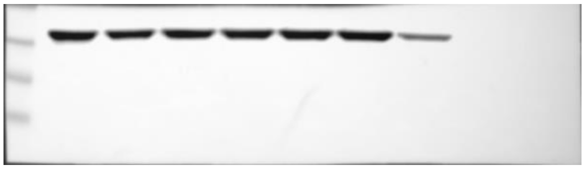

ACTB, NK-92

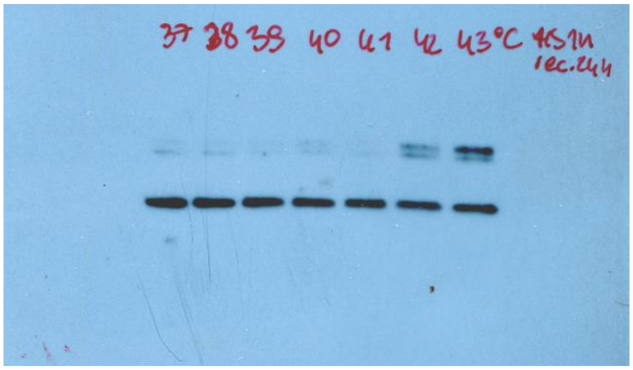


PDCD1, Jurkat

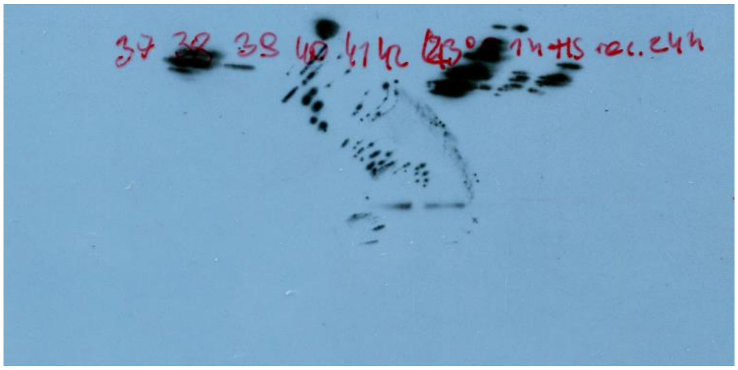

HSPA1, Jurkat

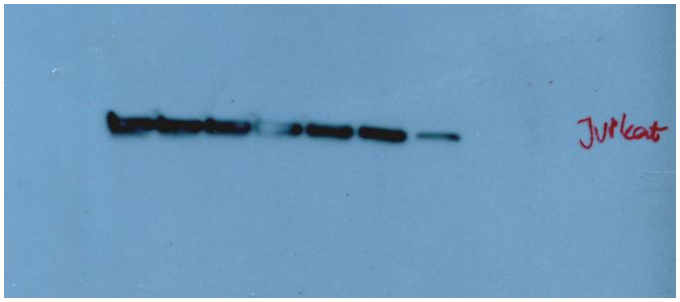


ACTB, Jurkat

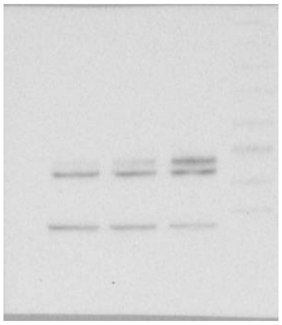

PDCD1, GM07062

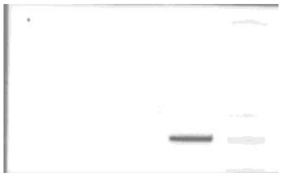

HSPA1, GM07062

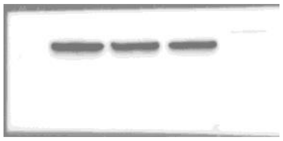

GAPDH, GM07062

**Figure S2E**

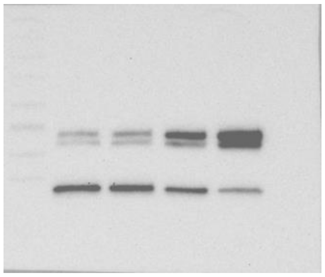

PDCD1, NK-92

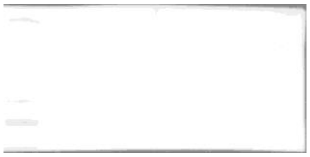

HSPA1, NK-92

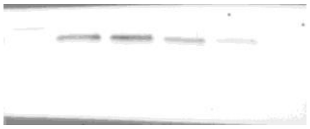

GAPDH, NK-92

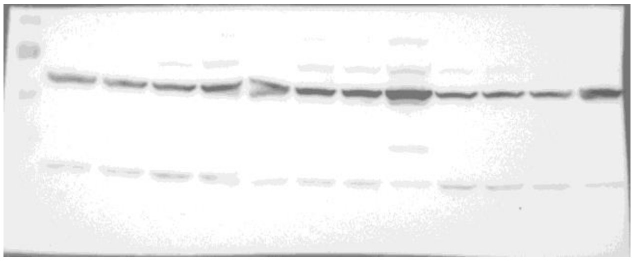

PDCD1, Jurkat, THP1, GM07062

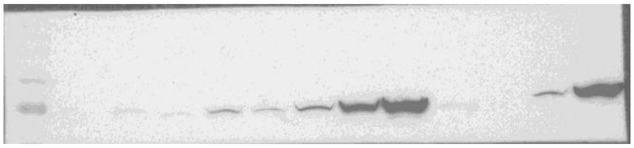

HSPA1, Jurkat, THP1, GM07062

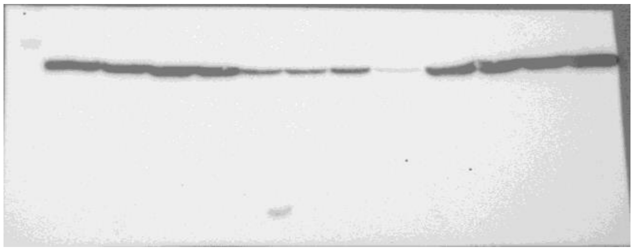

GAPDH, Jurkat, THP1, GM07062

**Figure S2F**

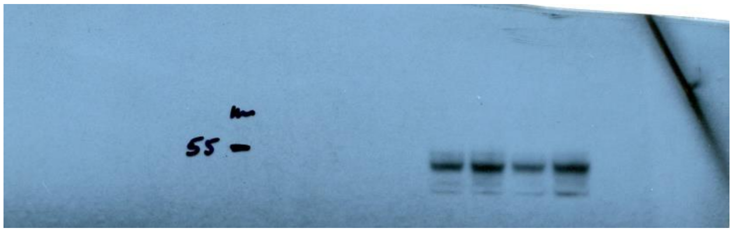


PDL1

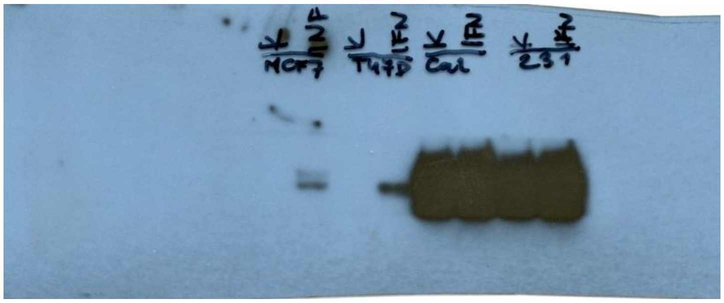

PDL1, overexposed

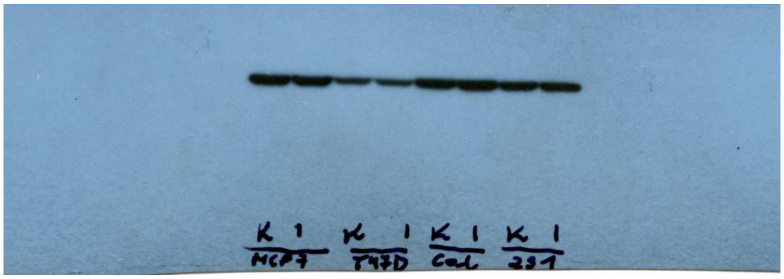

GAPDH
